# Structure and dynamics of the central lipid pool and protein components of the bacterial holo-translocon

**DOI:** 10.1101/490250

**Authors:** Remy Martin, Andreas Haahr Larsen, Robin Adam Corey, Søren Roi Midtgaard, Henrich Frielinghaus, Christiane Schaffitzel, Lise Arleth, Ian Collinson

## Abstract

The bacterial Sec translocon, SecYEG, associates with accessory proteins YidC and the SecDF-YajC subcomplex to form the bacterial holo-translocon (HTL). The HTL is a dynamic and flexible protein transport machine capable of coordinating protein secretion across the membrane, and efficient lateral insertion of nascent membrane proteins. It has been hypothesized that a central lipid core facilitates the controlled passage of membrane proteins into the bilayer, ensuring efficient formation of their native state. By performing small-angle neutron scattering (SANS) on protein solubilized in match-out deuterated detergent, we have been able to interrogate a ‘naked’ HTL complex, with the scattering contribution of the surrounding detergent micelle rendered invisible. Such an approach has allowed the confirmation of a lipid core within the HTL, which accommodates between 8 and 29 lipids. Coarse-grained molecular dynamics simulations of the HTL also demonstrate a dynamic, central pool of lipids. An opening at this lipid rich region between YidC and the SecY lateral gate may provide an exit gateway for newly synthesized, correctly oriented, membrane protein helices to emerge from the HTL.

## Introduction

The general process of protein secretion and membrane protein insertion is achieved by the conserved secretory, or Sec, machinery at the plasma membrane of bacteria and archaea, and the endoplasmic reticulum (ER) of eukaryotes. The protein-conducting channel is formed by a core hetero-trimeric assembly – the SecY of bacteria and archaea, and Sec61 complex of eukaryotes (1, 2) – through which secretory and membrane proteins are driven, respectively across and into the membrane. This process occurs either during protein synthesis, involving the direct binding of co-translating ribosomes to the Sec complex, or post-translationally, powered by associated energy transducing factors, such as the ATPase SecA in bacteria (3, 4).

Additional components combine with the core complex to facilitate the lateral passage of trans-membrane α-helices into the bilayer, or for the implementation of specific modifications, like glycosylation in eukaryotes. Indeed, the structure of the eukaryotic holo-translocon engaged with the ribosome illustrates how the core complex and accessory factors could streamline the efficient translocation and glycosylation of proteins at the ER membrane (5).

The bacterial core translocon SecYEG associates with the ancillary sub-complex SecDF-YajC (6) and YidC (7) to form a 7 protein super-complex *aka* the holo-translocon (HTL) (8). Generally, secretion through the translocon occurs post-translationally, whereas membrane protein insertion is co-translational (9). The HTL ensures efficient translocation, folding and assembly of secretory and membrane proteins and can be produced in sufficient quantities for structural and functional analyses (8, 10, 11). Its availability enabled a preliminary structural analysis combining electron cryo-microscopy (cryo-EM) and small-angle neutron scattering (SANS) (12) (Figure 1). Interestingly, the proteins are arranged around a central cavity, most likely constituted of lipids, which we proposed to form a protected environment for the co-translational insertion of trans-membrane α-helical bundles. The encapsulation of nascent unfolded membrane proteins would prevent catastrophic proteolysis or aggregation and thus promote efficient protein folding, much in the same way that GroEL facilitates the folding of globular proteins within a secluded hydrophilic chamber (13).

**Figure 1).**
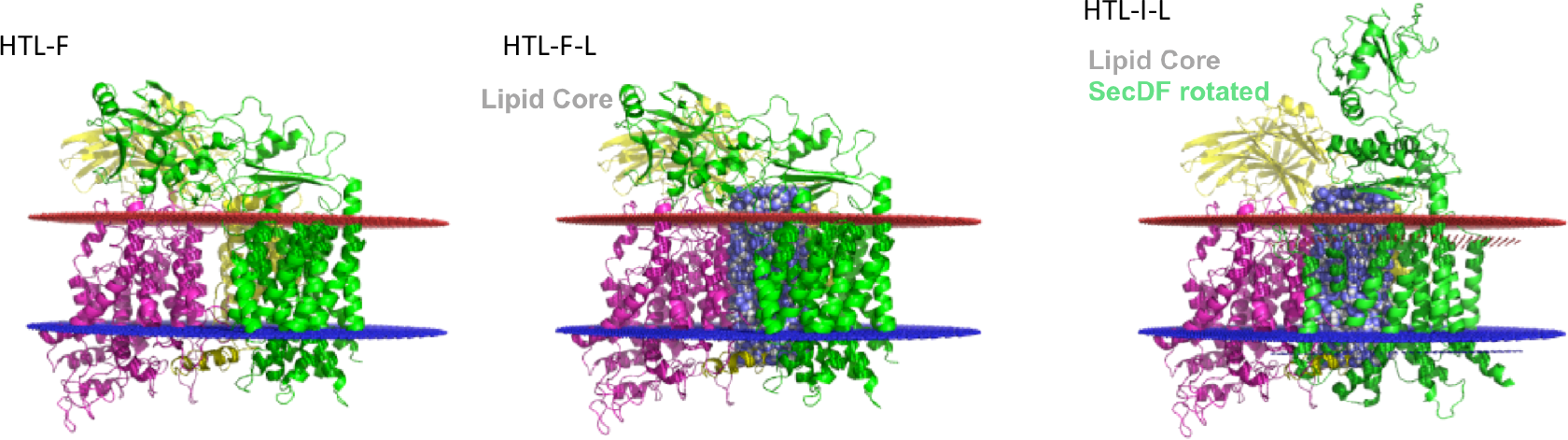
Atomistic models of HTL with and without a ‘hand’ positioned lipidic core. 3 HTL structures used to fit the experimental SANS data. HTL-F is the starting structure (HTL with SecDF in the F-form), and is based on the EM fitted structure from Botte et al. 2016. SecYEG is shown in magenta, with SecDF in green, and YidC in yellow. HTL-F-L is the same protein arrangement with the addition of a lipid core. HTL-I-L is the same structure as HTL-F-L (i.e. contains lipids) but has had the SecDF P1 domain rotated (HTL with SecDF in the I-form). Lipid bilayer planes are marked in red (periplasmic side) and blue (cytoplasmic side).

High-resolution structures of the individual components of the HTL are known (14–16), and they could be fitted into the low-resolution cryo-EM structure to create a preliminary atomic model of the HTL, supported also by biochemical data (12). In this model the lateral gate of SecY, through which nascent trans-membrane helices enter the membrane (14), faces the central lipid cavity. YidC is located on the opposite side of the cavity, with its putative binding site for inserting trans-membrane helices (15) also facing the lipid pool. The juxtaposition of these regions at the proposed central lipid core of the HTL provides a compelling case for their concerted action in membrane protein insertion.

To explore further the structure and arrangement of the central lipid pool we conducted an analysis of the HTL, combining SANS and coarse-grained (CG) molecular dynamics (MD) simulations. The HTL was solubilised in matched-out deuterated detergent n-dodecyl-β-D-maltoside (d-DDM). This d-DDM was deuterated separately in the head and tail group to fully match out the neutron contrast of the detergent in a 100% D_2_O-based buffer (17). This way the detergents become invisible in the SANS experiment. This allowed us to distinguish and describe the lipid component of the translocon. The CG MD simulations support the notion of a stable and persistent lipid-filled cavity within the centre of the HTL. Beyond this we discuss the role of such a lipid pool in the insertion and folding of membrane proteins *via* the Sec machinery.

## Materials and Methods

### HTL preparation and d-DDM exchange

HTL was purified as described previously (8). Purified HTL in hydrogenated DDM was exchanged into a 100% D_2_O buffer containing deuterated DDM. Detergent exchange was performed on a Superose 6 (10/300) column equilibrated in a simple TS buffer (20mM Tris pH 7.5, 100mM NaCl_2_), made with 100% D_2_O, and 0.02% deuterated DDM.

### SANS data collection for deuterated detergent

Samples were prepared and measured in 2 mm quartz cuvettes (Hellma), temperature controlled at 10 °C. SANS data were collected on KWS-1 at FRMII at MLZ (Garching), at a wavelength of *λ* = 5 Å and a wavelength spread of *Δλ/λ* = 10% (FWHM). Sample-detector/collimation distances of 1.5m/4m and 8m/8m were used, to obtain a *q*-range of 0.006 to 0.44 Å^-1^, with a good overlap between the settings. The wave vector, *q*, has the usual magnitude of 4*π* sin(*θ*)/*λ*, where 2*θ* is the scattering angle. Transmissions were measured at a 4m/4m setting with 3 min exposure time. Data were calibrated using plexiglass as a calibrant to yield the absolute scaled scattering intensity, *I*(*q*), in units of cm^-1^.

Correction and averaging was performed using QtiKWS (v. 10; www.qtikws.de), and the buffer measurement was subtracted subsequently. The sample was measured for ~4 hours (1.75 hours at the 8m/8m setting and 2 hours at the 1.5m/4m setting) to obtain sufficient signal over background. 15 minute measurement windows were used to monitor change in scattering over time. No change was observed, meaning that the sample was stable during the measurements.

### SANS data analysis

A combination of the home-written software CaPP (v. 3.9; github.com/Niels-Bohr-Institute-XNS-StructBiophys/CaPP) and WillItFit (18) was used to fit the data. A 3 Å thick water layer with 6% higher scattering length than bulk D_2_O was added (19) but was excluded from the transmembrane region. A hydrophobic bilayer thickness of 30.6 Å was assumed in accordance with the orientations of proteins in the OPM membrane database (20). Resolution effects were included using the resolution width, *Δq*(*q*), present in the 4^th^ column of the data files provided by the beamline. CaPP was also used to calculate the theoretical pair distance distribution functions, *p*(*r*), for the atomistic structures. Experimental *p*(*r*) were calculated using BayesApp (21) including a constant background in the fit and truncation of data at q = 0.3 Å^-1^. The fit to obtain the *p*(*r*) had a 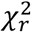 value of 2.7. As the model is generic and thus true for this dataset as well, this value was expected to be close to unity. There were 112 data points in the fitted range, and the degrees of freedom of the model was estimated as *N*_*g*_ = 8.3 (Vestergaard & Hansen 2006). The probability of obtaining a 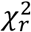 of 2.7 given 112 points and *N*_*g*_ = 8.3 is only ~10^-16^ for a true model. We could therefore conclude that the experimental errors were underestimated and they were renormalized by *σ*_*new*_ = β · σ_*old*_, where 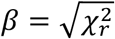

The forward scattering, as determined by Guinier analysis (Figure S1), can be used to calculate a model-free estimation of the number of lipids in the lipid core. The protein concentration of the sample was calculated from a measurement of the UV280 absorption of 0.65 cm^-1^, and an extinction coefficient of 234600 cm^-1^ M^-1^, as calculated from the protein sequence, using ExPASy ProtParam (web.expasy.org/protparam). The forward scattering from the protein-lipid complex (HTL + lipid core) is given as:

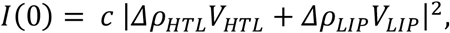

where *c* is the concentration (number of complexes per cm^3^), *Δρ*_*HTL*_ and *Δρ*_*LIP*_ are the excess scattering length densities (scattering contrasts) of the protein and lipid respectively, and *V*_*HTL*_ and *V*_*LIP*_ are the corresponding volumes. The sample was purified with an *E. coli* lipid extract (avantilipids.com/product/100600), with known lipid composition, so *Δρ*_*LIP*_ could be estimated. The only unknown was therefore *V*_*LIP*_, the volume of the lipid core:

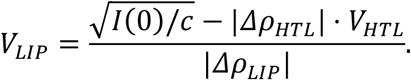

*V*_*LIP*_ is calculated by subtraction of two numbers, 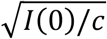 and (|*Δρ*_*HTL*_ | · *V*_*HTL*_), equal in magnitude, and each with an associated uncertainty, which result in a relatively large error on the calculated result. The major contributions to the uncertainty stems from the absorption measurement used to estimate the molar concentration. We assumed a 15% uncertainty on the concentration measurement, 10% on the estimation of *I*(0) and 2% on the estimated volumes of HTL and the lipids. The number of lipids could then be found by dividing *V*_*LIP*_ by the mean volume of the *E. coli* lipids (1216 Å^3^), which was calculated from the lipid composition (avantilipids.com/product/100600) using known volume for the different lipid components (22).

A fit was made where the fitting algorithm was allowed to mix HTL-F-L and HTL-I-L (Figure 1) to obtain the optimal fit (Figure 2). The intensity of the mix was given as:

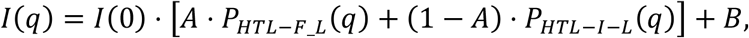

where *A* is the fraction of the sample in the HTL-F-L form.

**Figure 2).**
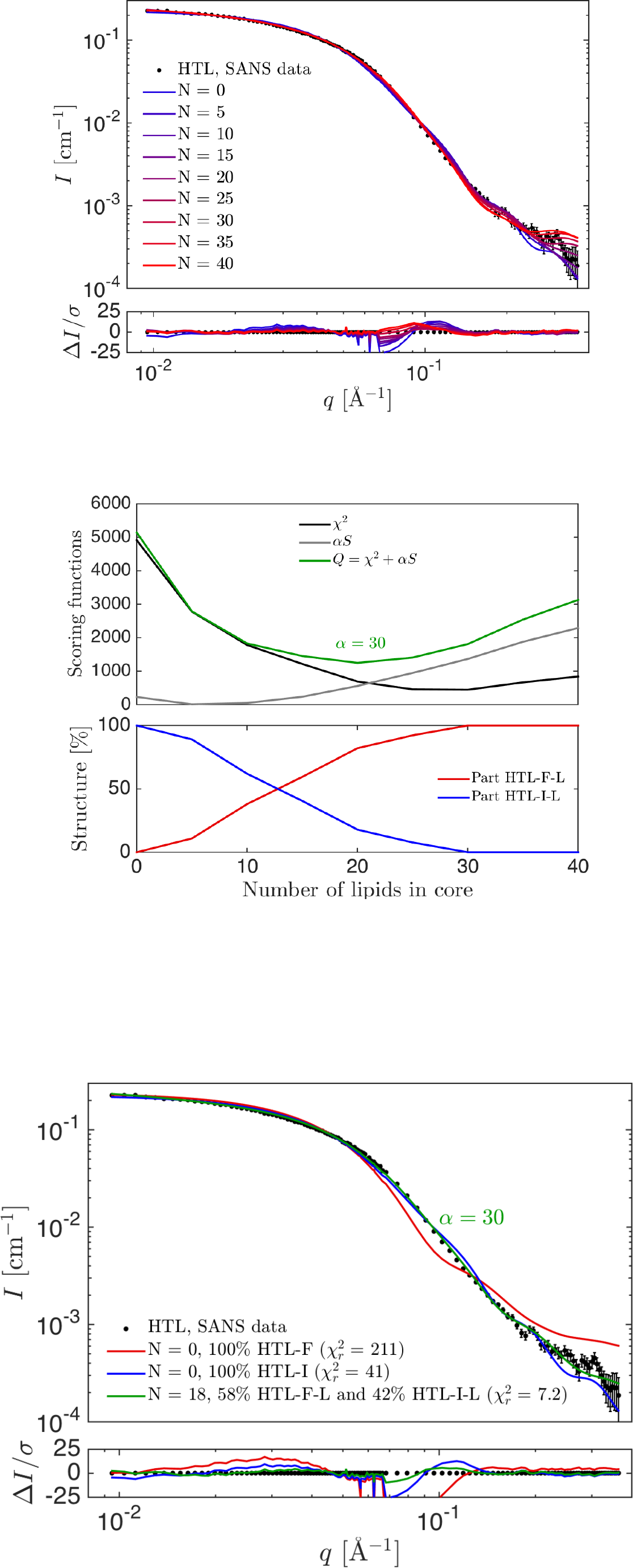
Model fitting of theoretical scattering to experimental SANS data of HTL in d-DDM. A) Theoretical scattering of a linear combination of model HTL-F-L and HTL-I-L in which the number of central lipids is varied from 0-40 (red-blue), plotted against experimental HTL data (black dots). B) Upper panel shows χ^2^, *αS* and *Q* (see text) for the fits and lower panel shows the amount of HTL-F-L and HTL-I-L in the fits. C) Fit to data with HTL-F (no lipids) in blue, HTL-I (no lipids) in blue, and the most probable fit for *α* = 30 (see text), with 18 lipids in the core, 58% in the HTL-F-L form and 42% in the HTL-I-L form, and with ~1% of the total protein being aggregated.

The goodness of the fits was evaluated using the reduced *χ*^2^, given as 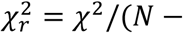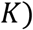, where *N* is the number of datapoints and *K* the number of fitting parameters. The *χ*^2^ is defined in terms of the measured experimental intensities 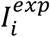 and corresponding uncertainties *σ*_*i*_ and the fitted theoretical intensities 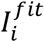:

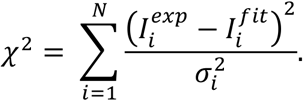

There was a minor contribution of aggregates in the sample, as seen from the upturn in the Guinier plot (SI, Figure S1). The presence of aggregations were also clear from the “tail” of the *p*(*r*) with a large maximal interparticular distance, D_max_, of ~200 Å (Figure 3). The aggregate contribution was taken into account in the fits (Figure 2) by including a fractal structure factor, *S*_*frac*_ (*q*), to the model, as previously described (23). Shortly, a fractal aggregate description was used (24) in combination with the decoupling approximation (25) and the form factor of the complex, *P*(*q*):

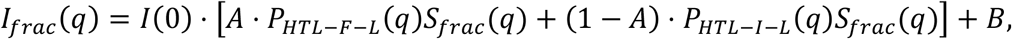

where *S*_*frac*_ (*q*) is the effective form factor after the decoupling approximation was applied. A mean radius of *R* = 42.1 Å was used for HTL, corresponding to the radius of a sphere with volume equal to the sum of Van der Waals volumes of the atoms in the protein (26). The models were implemented in WillItFit (18).

**Figure 3).**
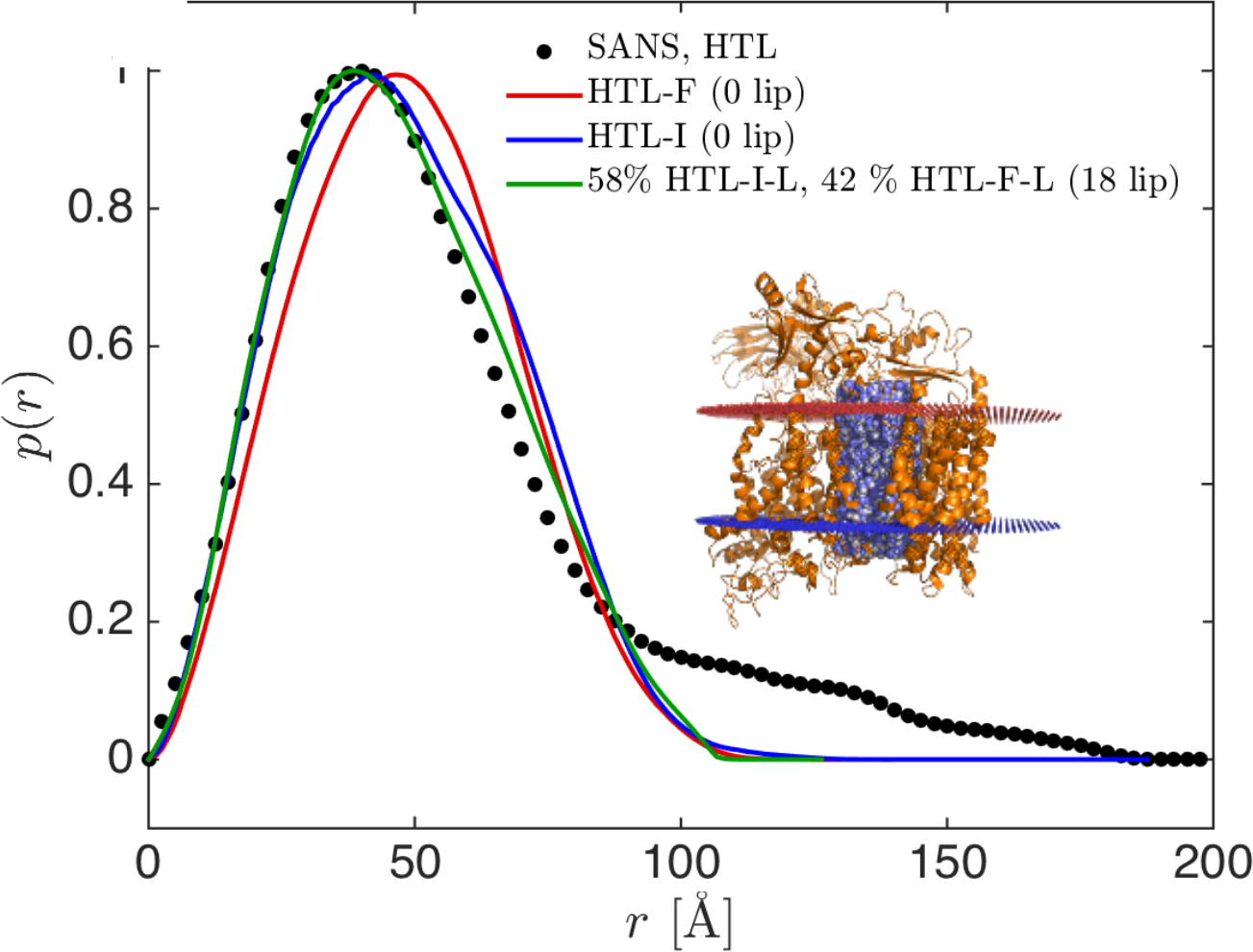
Pair distance distribution functions *p*(*r*) for experimental SANS data of HTL. *p*(*r*) plot of HTL SANS data (black), HTL-F (no lipids, red), HTL-I (no lipids, blue) and the Fourier transform of the most probable fit for α = 30 (see text), with 18 lipids, and 58% of HTL in the F-form and 42% in the I-form (green). The inset image shows the HTL-F-L in cartoon representation (orange) with a lipid core representative of 18 lipids (blue and white). Lipid bilayer planes are marked in red (periplasmic side) and blue (cytoplasmic side).

### Molecular Dynamics

CG MD simulations were built according to the MemProtMD protocol (27), using PDB 5MG3 as an input. Simulations were run for 350 ns with elastic network restraints of 500 kJ mol^-1^ mn^-2^ between all protein beads within a cut-off distance of 1 nm, at 310 K using 20 fs time steps. These simulations were then extended to 3 μs with elastic networks only applied to beads within 1 nm and on the same protein chain. This was done to allow the central lipid pore to change size without the restriction of inter-subunit elastic networks. Post simulation snapshots were converted to atomistic description (28) for comparison with the SANS data.

Simulations were run on Phase 3 of BlueCrystal, the University of Bristol’s High Performance Computer (HPC). Images of proteins were made in PyMOL and VMD, and data were plotted with gnuplot or matplotlib.

## Results

### SANS confirms a central lipid core within the HTL

Previous studies show that the purified HTL complex is composed of its constituent protein subunits and significant proportions of lipid and detergent (12). In the purified complex, the majority of this lipid and/ or detergent component is localised at the centre of the complex with the protein at the periphery. Due to the relatively close contrast match points of DDM (21.7% D_2_O) and *E. coli* lipids (13.1% D_2_O) it is difficult to distinguish and separate the scattering contributions from the lipid and the solubilizing detergents, and therefore identify whether the central scattering contribution was attributable to lipid or detergent. In order to address this, SANS experiments were performed on the HTL using partially deuterated DDM (d-DDM) to mask the scattering signal associated with the detergent. The DDM sugar head group and tail moieties were chemically deuterated independently, with differing levels of deuteration, such that the scattering length densities of both head and tail group of the d-DDM are equivalent to that of 100% D_2_O.

Purified HTL was detergent exchanged into d-DDM buffer by gel filtration chromatography (see Materials & Methods), wherein the d-DDM buffer was made up with 100% D_2_O. So, the recorded measurements were conducted at the d-DDM contrast match point and with minimum incoherent scattering background from the buffer. Thus, only scattering contributions of the protein and lipid components were measured and the detergent rendered effectively invisible.

Guinier analysis of the collected data indicates a radius of gyration (*R*_*g*_) for the HTL in absence of the DDM scattering contribution as 41.1 ± 0.3 Å, slightly higher than the calculated theoretical *R*_*g*_ of 37.0 Å. The forward scattering, *I*(0), determined by Guinier analysis (29), can be used to calculate a model-free estimation of the lipid volume of the HTL (see Materials & Methods). From an *I*(0) value of 0.23 cm^-1^, the volume of the lipid pool (*V*_*LIP*_) can be estimated to 12000 Å^3^. Assuming a mean volume of an *E. coli* lipid as 1200 Å^3^ (calculated from (22)), the model-free estimation indicates the presence of a lipid pool consisting of 10 lipids, supporting a significant lipid-based scattering contribution.

However, due to the nature of the determination of *I*(0) and the cumulative uncertainties involved in calculating *V*_*LIP*_, the estimated error on the result was +/−17, as obtained by error propagation. Moreover, the aggregation of the sample, as evident from the upturn in the Guinier plot (Figure S1) and the “tail” of the pair distance distribution function (Figure 3), would result in an overestimated value for the forward scattering, and thus of the estimated number of lipids. A more precise estimation could be obtained by model-based analysis, where the full q-range was investigated and the aggregates could be included in the model.

A model lipid core of this volume was created, with the height corresponding to a typical lipid bilayer (50 Å). A simple cylindrical shape was assumed for the lipid core (Figure 1), which spanned the height of the trans-membrane part of HTL. The lipid core consisted of beads homogeneously filling the cylindrical volume with an average scattering length density corresponding to the average scattering length density of the lipid extract used in the sample purification and preparation. The cylindrical volume was filled with beads, each represented a dummy CH_2_ groups. The beads were placed in the cylindrical volume by a Monte Carlo approach. The lipid core was positioned in the central cavity of the preliminary EM-fitted HTL structure (PDB: 5MG3; (12)). The structure without lipids was termed HTL-F (HTL in the F form) and the structure with lipids HTL-F-L (Figure 1).

### Probing the flexibility of SecDF Periplasmic domains by SANS

In the SecDF-YajC sub-complex, the SecD periplasmic domain 1 (P1), has been observed in two distinct orientations, the F and the I form, with an approximate 100° rotation of P1 between the two structures (16, 30). To assess the effect of domain flexibility, a model was created, taking the lipid containing HTL-F-L structure and replacing SecDF in the F form with SecDF in the I form (PDB: 5XAM). This model was termed HTL-I-L (Figure 1) and utilized in subsequent data fitting process below.

### Refining the number of lipid and the structural flexibility

Theoretical scattering was calculated for a series of HTL-F-L and HTL-I-L structures containing a varying number of lipid molecules in the core (0-40) to find the most probable model (Figure 2A). The model fit to the experimental data improved as the number of lipids increases up to a count of 29 lipids, as assessed by the calculated 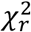 values (Materials & Methods), and worsens at numbers above this value (Figure 2B). However, the data were fitted on absolute scale with a scaling parameter, *k*, for the forward scattering. Optimally, *k* should be unity, and deviation from unity indicates a less probable model. But some deviation from unity was expected due to the uncertainty of the estimation of *I*(0) (see Materials and Methods). We therefore introduced a penalty function *S*(*k*):

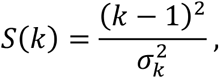

that increases as *k* differs from unity. 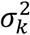 is the estimated uncertainty of *k*, which was set to 0.05. The most probable solution could then be found where the function *Q* = χ^2^ + *αS* was minimized (31), where *α* determines the weight given to *χ*^2^ and *S* respectively. *Q*, *χ*^2^ and *αS* are plotted on Figure 2B for *α* = 30. With this regularized expression, a lower number of 8 bound lipids were estimated, as both *S* and *χ*^2^ increase as *N* decreases below 8. Similarly, an upper limit of 29 lipids was determined, as both scoring functions increase as *N* rises above this value.

The optimal value depends on the choice of *α*, which is not trivial to determine, and which we will not go into detail here, but details can be found in Larsen et al. 2018 (31). In Figure 2C, the best fit for *α* = 30 is given, with *N* = 18, in the middle of the ‘allowed’ range. Of the whole range (between 8 and 29 lipids), the best fit is a linear combination of the HTL-F-L and the HTL-I-L structures; *e.g.* for *N* = 18, the model that best fits data has 58% of HTL-F-L and 42% of HTL-I-L. As lipid numbers increase, the calculated proportion of HTL-I-L increases as well. The number of lipids and the structural conformation of HTL are thus highly correlated parameters, but the most probable model has some structural flexibility of the domain, in addition to a significant scattering contribution from a lipid core.

All models included a portion of aggregates, varying from below 1% for the model with 8 lipids and above 5% for the model with 29 lipids in the core. The model with 18 lipids in the core included around 1% of aggregates.

In summary, the SANS analysis suggests that HTL has a lipid core of between 8 and 29 lipids and exhibits some flexibility of the periplasmic part of the SecDF domain.

### Coarse-grained simulation supports the existence of a lipid core

In order to assess the stability of the HTL complex, and begin characterisation of a central lipid core to the complex as indicated by SANS, a coarse-grained (CG) molecular dynamics (MD) study was performed. An atomic model of HTL was constructed using *E. coli* YidC (32), SecYEG (33), and *E. coli* homology models from *T. thermophilus* SecDF (12, 34). These structures were arranged to fit the experimental cryo-EM density of the HTL (PDB: 5MG3). The atomic structures were converted to CG models using the Martini forcefield (35, 36), and inserted into a simulation box filled with randomly oriented CG lipids (65% POPE, 30% POPG, 5% cardiolipin), and solvated with CG water and ions. The system was allowed to self-assemble, forming a clear lipid bilayer around the HTL. Following bilayer formation, and an initial settling period the radius of gyration of the protein complex settles at approximately 38 Å (Figure 4), in excellent agreement with the scattering data (~ 40 Å). The HTL was simulated for a total duration of 3 μs is (Figure 4B) and remained stable as determined using RMSD and *R*_*g*_ analysis (Figure 4C).

**Figure 4).**
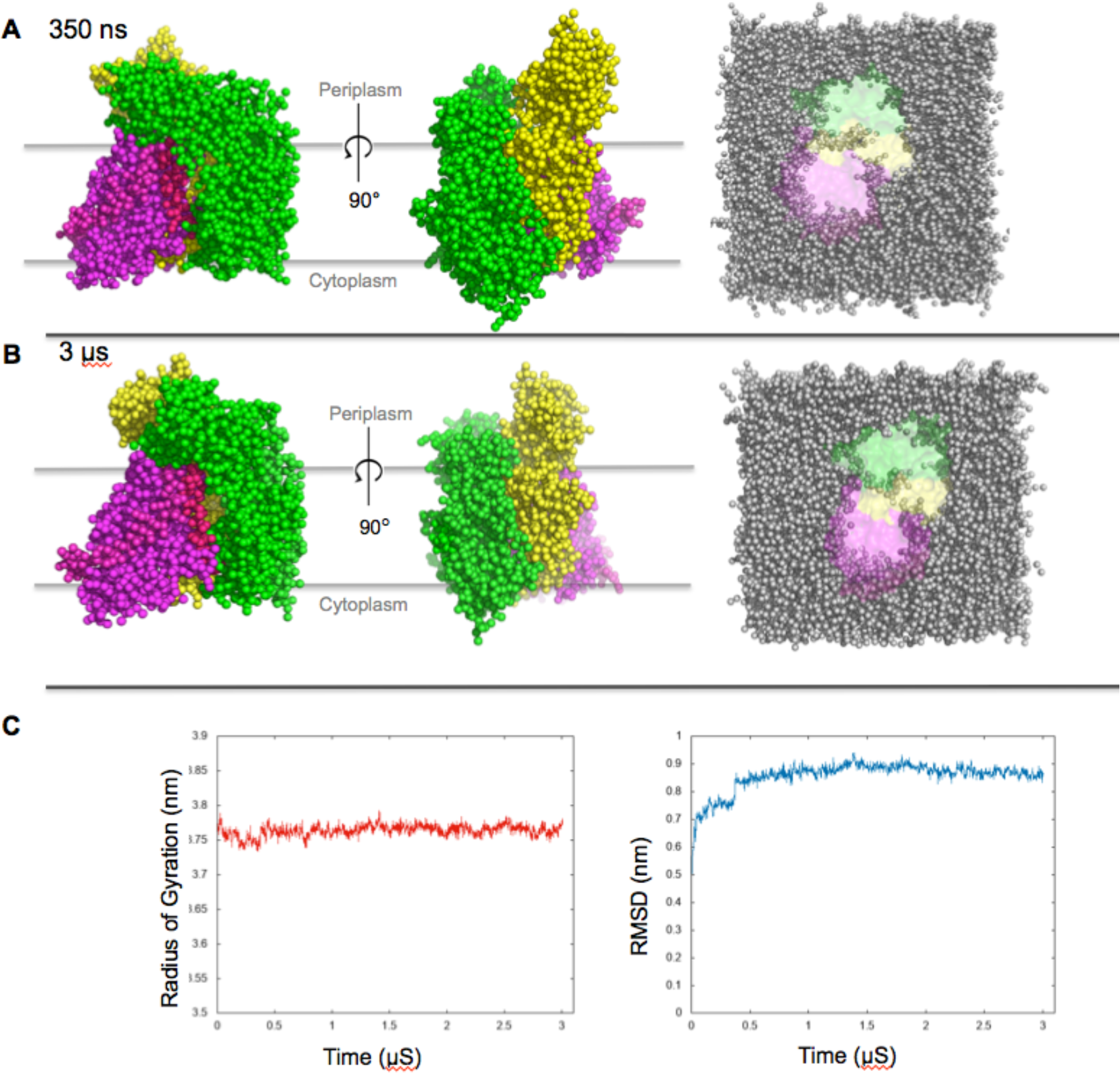
Coarse-grained HTL model, pre- and post-simulation. Coarse-grained HTL after 350 ns and 3 μs simulation. A) HTL shown after 350 ns simulation, viewed transversely through the membrane from two orientations, and from cytoplasmic face, showing the lipid arrangement within the complex. SecYEG is shown in Magenta, with SecDF in green, and YidC in yellow. B) As previous, but after 3 μs of simulation. C) Graphs showing the stability of the structure over 3μS, from l-r: Radius of gyration, RMSD.

CG modelling of the HTL complex shows the presence of a stable lipid pool at the interface between the trans-membrane domains of all of the components of the HTL, at the centre of the complex (Figure 5). The number of lipids within this island remains between 7 and 13 for the entire 3 μs duration of the simulation (Figure 5B). The average number of lipids remaining in the centre of the HTL complex is 9.4 +/- 0. 8 lipids, for the final 2 μs of a 3 μs simulation. Lipids are seen to diffuse in and out of the pool, predominantly through the gap between the SecY lateral gate and YidC, which may act as an opening point of the complex. Due to lipid diffusion, the lipid pool fluctuates in shape throughout the simulation, but remains between approximately 20 and 40 Å in diameter depending on the number of lipids present.

**Figure 5).**
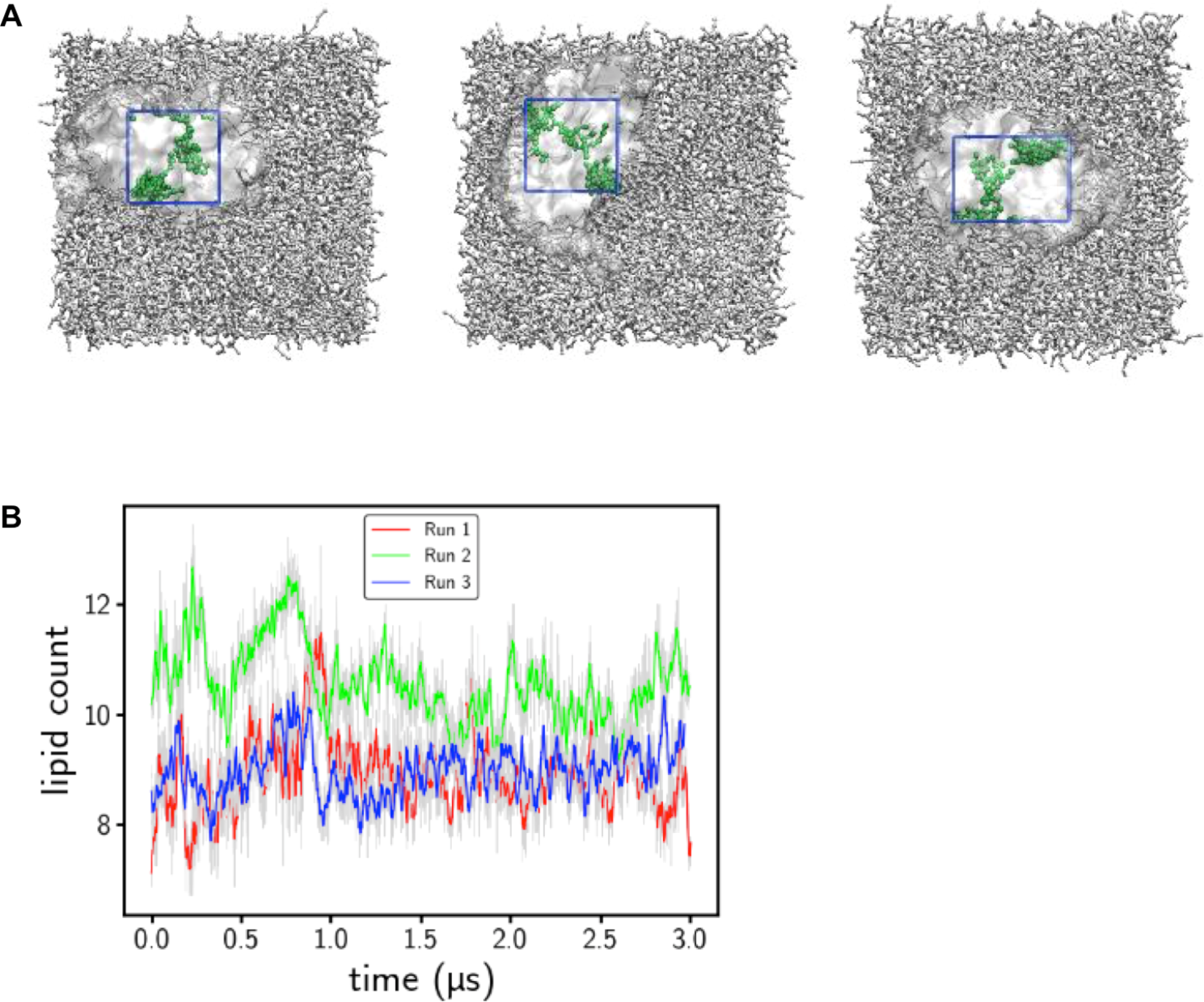
Localisation and number of lipids within the HTL during MD. The localisation of the lipids within the HTL during simulations. A) Shots of 3 independent coarse-grained simulations of HTL in a mixed lipid bilayer. In each image, the lipids present in the centre after 3μs simulation are highlighted green. A boundary box was created for each simulation, and lipid presence within the area quantified. B) Graph showing the number of lipids within the core of the HTL, as defined by the boundary box, over the course of the simulation time.

A CG HTL model was extracted from the last frame of the simulation, and converted to atomistic model with 7 POPE and 2 POPG in the core. This model had an overall structure similar to HTL-F-L. A model of HTL-I-L with the lipids from the simulations was also generated. The scattering from these models were calculated and compared with data (Figure 6). The HTL-I-L with the core of CG MD lipids fitted the data quite well 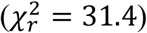, whereas the HTL-F-L with CG lipids resulted in a poor fit 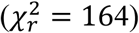. Less than 1% of aggregates were included in the two fits.

**Figure 6).**
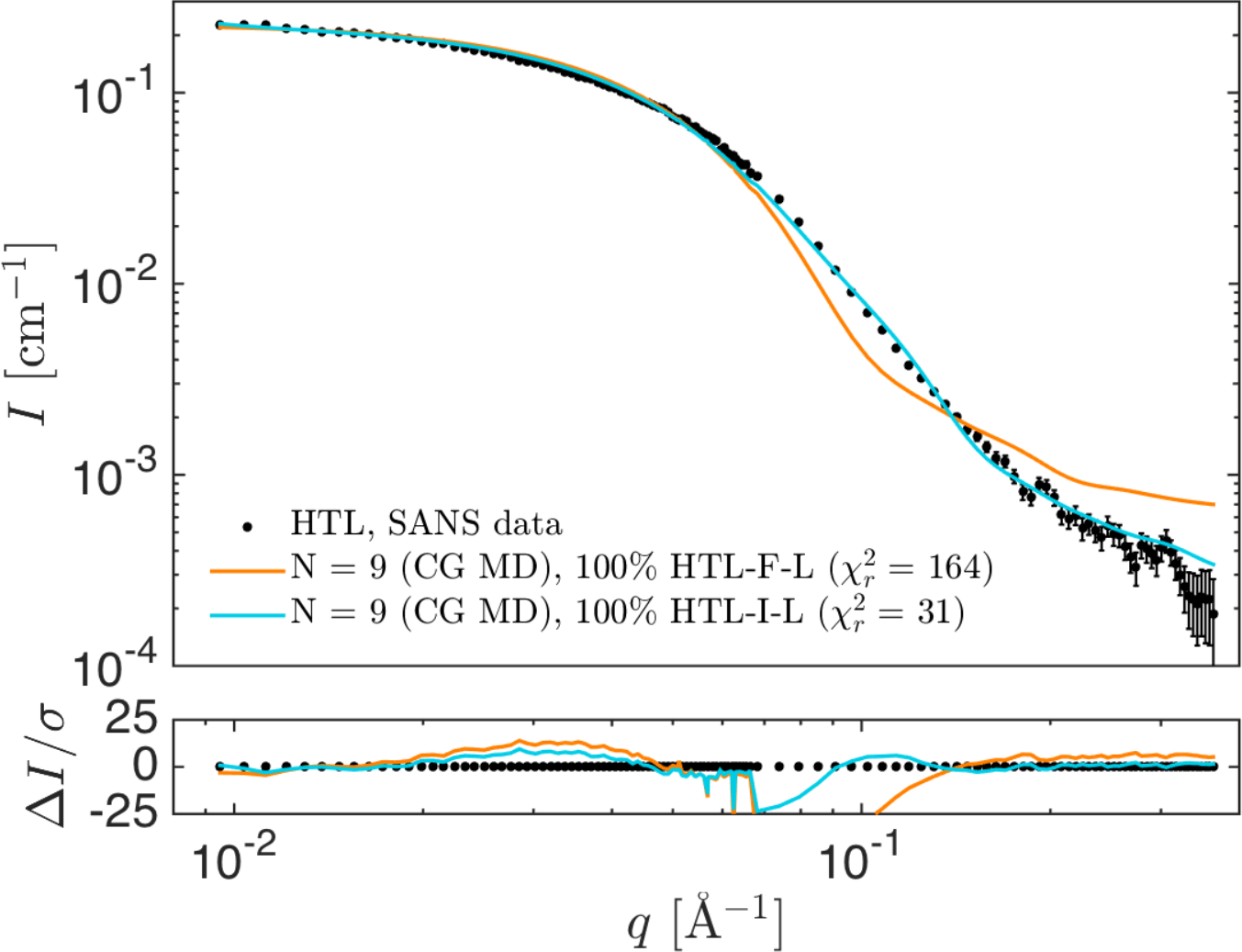
Model fit of HTL with post-simulated lipids to experimental SANS data. A) Experimental HTL SANS data (black dots) fitted with the HTL-F-L structure with the 9 lipids from the CG MD simulation (orange) and with the HTL-I-L structure with the 9 CG MD lipids (cyan).

## Discussion

The results presented show the HTL to be a dynamic complex, unequivocally demonstrating that the individual subunits are arranged around a central lipid core. The SANS data supports a model of the HTL containing a pool of between 8 and 29 lipids at its centre. The fit is improved accounting for flexibility of SecDF indicating that a significant part of the proteins have a rotated SecD periplasmic domain.

The lipid pool at the centre of the HTL complex was observed to be stable during the CG MD simulations. This correlates well with both the SANS data in this study, and previous structural studies of the HTL, indicating protein is located towards the periphery of the particle in solution with lipid and/ or detergent material located towards the centre (12). The number of lipids observed in the central core during the simulation remained between 7 and 13 (Figure 5), towards the lower range of the estimate provided by SANS. The lipids were observed to diffuse in and out of the core during the simulations, suggesting that there is natural fluctuation of the lipid core volume in bilayer conditions.

In detergent solubilized conditions, it has been shown that prolonged exposure to detergent can remove associated annular lipids from membrane proteins (37, 38), though the integral lipids within the centre of the HTL are not likely to be as easily dissociated. However it is feasible that DDM exposure may destabilize the HTL, potentially facilitating exchange of some central lipids into surrounding detergent micelles, raising the possibility that the lipid numbers estimated by the SANS models may be lower than the true physiological value.

There was difference between the applied lipid cores obtained by MD simulations and used in the SANS analysis. The lipid core used in the SANS analysis, and which fitted the data best, was a small cylindrical bilayer restrained to the centre of the complex and spanning the entire TMD. The lipids in the CG MD simulation on the other hand formed a monolayer that penetrated the complex in the bilayer plane (Figure 7). None of the two lipid core models are perfect descriptions. The cylindrical model is very simplified, with a homogeneous scattering contrast, a fully symmetric morphology. Moreover, the model was not checked for steric clashes between lipid and protein and should therefore not be considered a fully physical model. For that reason, it is not surprising that even the most probable models did not fit the data perfectly (Figure 2C). Some systematic discrepancies were apparent, especially around 0.1 Å^-1^ (as clearly seen in the residual plot) and a 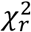 of 7.2 with 18 lipids in the core; *i.e.* still relatively far from unity.

**Figure 7).**
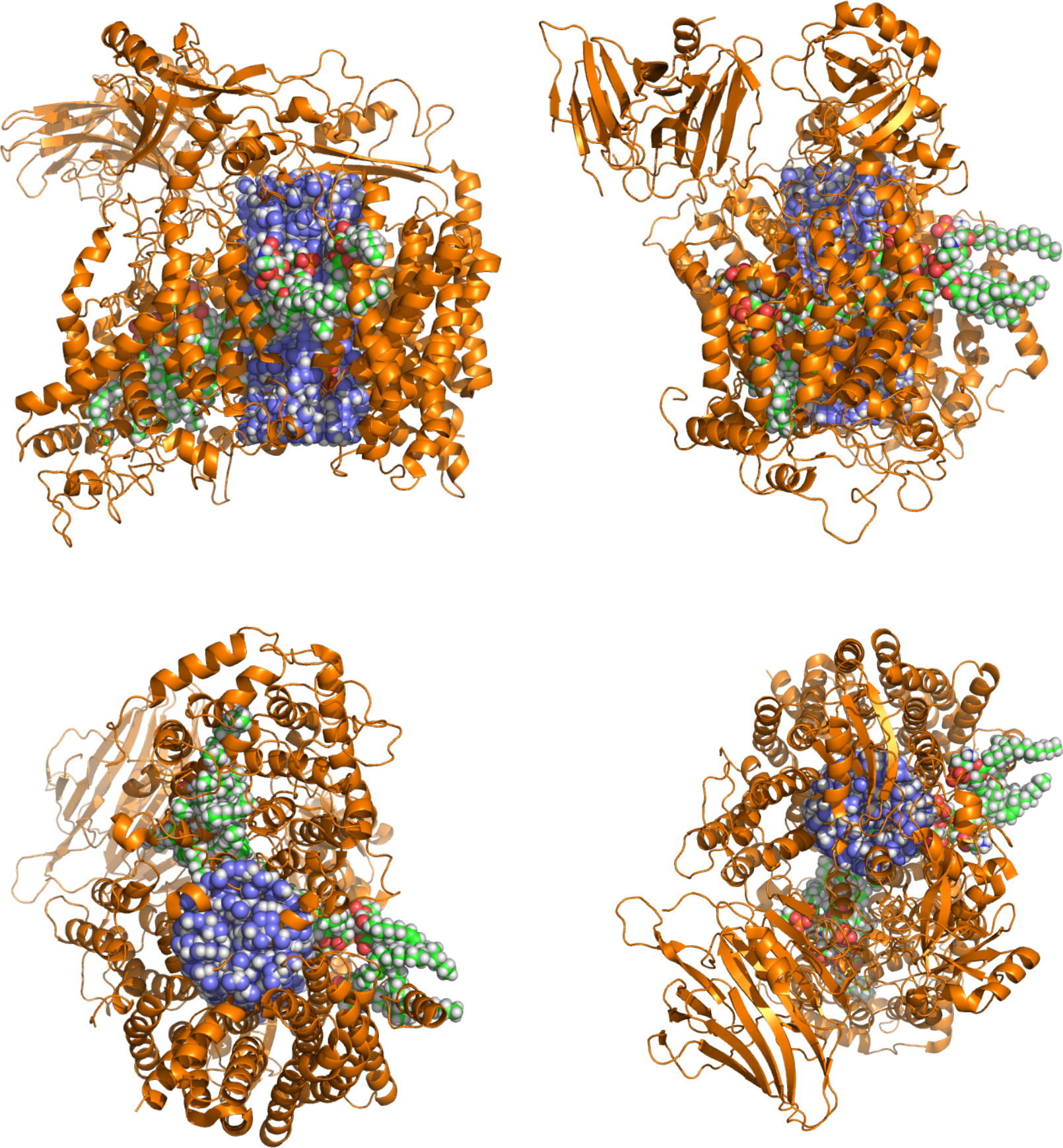
Atomistic comparison of HTL with ‘hand’ positioned and simulated lipid core. The HTL complex (PDB: 5MG3, orange) with two different models for the lipid core. The simple bilayer model used to fit the SANS data (blue/white, corresponding to 18 lipids), and the monolayer of lipids from the CG MD simulations (red/green/white, 9 lipids). Clockwise from top left: Front view, side view, top view, and bottom view.

The CG lipid model is more physical, as it is contrained with force fields, and has no steric clashes. The lipids are also better models for real lipids than a simple, homogeneous cylindrical lipid core. However, the CG lipid model did not fit the data as well as the simpler cylindrical model (Figure 6), as reflected in the 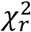 value of 31 for the best fit with HTL in the I-form. So, this data could possibly be used in future studies to benchmark and improve the applied force fields used for such challenging protein-lipid systems.

An ensemble of structures may well be a better representation for the dynamic complex than a single static structure, due to the diffusion of the lipids, and structural dynamic of the protein periplasmic parts. However, both lipid models, in their present form, provide valuable complementary information about the HTL proteo-lipid complex.

The SecD P1 periplasmic domain remained in close proximity to the other parts of the complex in the CG MD simulations, and may prevent formation of a periplasmic pool of lipids. SecD interacts with unfolded proteins (16), suggesting it may play a facilitative role in the HTL of binding and facilitating passage of periplasmic domains of membrane proteins as they are inserted by HTL. Insertion of a multi-spanning membrane protein by HTL may sequentially open up the complex as additional transmembrane helices insert, allowing lipids to fill the periplasmic side of the core.

The MD simulation points to a potential gateway in the cytoplasmic membrane face between SecY and YidC, through which lipids are able to diffuse in and out of the lipid pore (Figure 5). This gateway is capped at both ends by SecY residues previously identified as acidic lipid contact sites (39). During insertion, YidC is known to function in concert with SecY performing chaperone activities which facilitate correct folding of trans-membrane helices as they sequentially exit the lateral gate of SecY (40, 41). In this context the encapsulated lipidic microenvironment between this lateral gate and YidC would prevent aggregation and help achieve the native state of membrane proteins during the co-translational insertion process. This mechanism could operate for assisted folding of small membrane proteins, or for successively emerging helical bundles of larger polytopic substrates. The subsequent release of membrane proteins, or sequential release of their domains from the holo-complex would then be facilitated by the observed flexibility of the HTL (12), presumably by the partitioning of SecYEG and SecDF-YajC subcomplexes. This deployment of lipids for encapsulated membrane protein folding and controlled release presumably offers an effective means to efficiently achieve native states of monomeric proteins and assembly competent states for multi-subunit complexes.

## Acknowledgements

R.M. received a University of Bristol Postgraduate Scholarship. Additional support was gratefully received by I.C. from the BBSRC (BB/M003604/1 and BB/I008675/1). L.A. and A.H.L thank CoNeXT and University of Copenhagen for co-funding the project. The authors gratefully acknowledge the financial support provided by JCNS to perform the neutron scattering measurements at the Heinz Maier-Leibnitz Zentrum (MLZ), Garching, Germany. Part of this work is based upon experiments performed at the KWS-1 instrument and the authors would like to thank for the awarded beamtime. We would also thank Frederik Tidemand and Nicolai T. Johansen for great support during the SANS beamtime. We thank for the deuterated DDM (d-DDM) that was synthesised by Dr. Tamim Darwish (ANSTO, NSW, Australia).

## Author Contribution

R.M. and I.C. designed protein preparations for SANS, executed by R.M., R.M., C.S. and L.A. conceived the SANS experiments. R.M., S.R.M., A.H.L. and L.A. designed the SANS experiment, R.M., S.R.M., A.H.L. and H.F. conducted the SANS experiments and performed the initial analysis. A.H.L. and L.A. designed the SANS data analysis, which was then carried out by A.H.L. with inputs from L.A., R.M. and S.R.M., R.M. and R.A.C. designed the molecular dynamics experiments. R.A.C. performed the molecular dynamics experiments. R.M. and R.A.C. performed analysis of the molecular dynamics data. R.M., A.H.L. and IC co-wrote the manuscript with inputs from all co-authors.

## Supplementary figures

**Figure S1).**
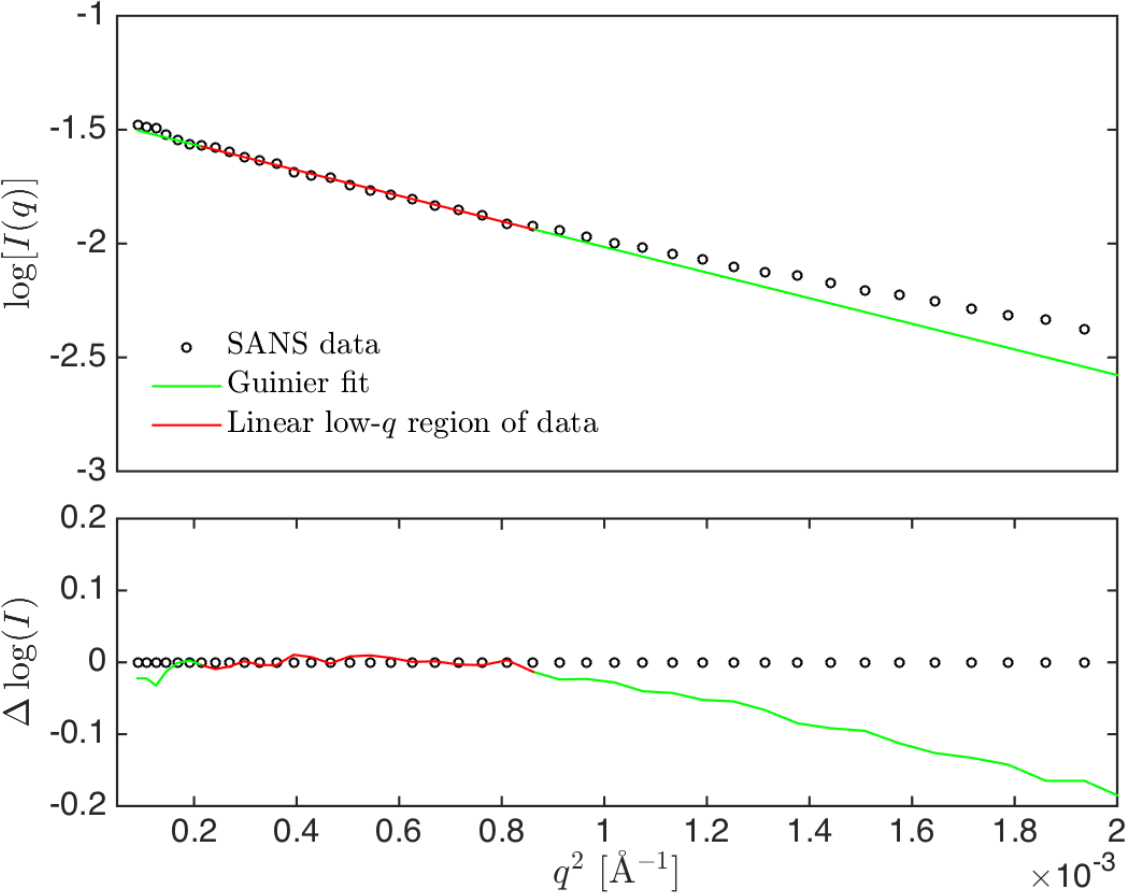
Guinier analysis of the SANS data of HTL. *R*_*g*_ was determined to be 41.1 +/- 0.3 Å, with *q*_*max*_*R*_*g*_ = 1.21. *I*(0) was determined to be 0.234+/-0.001 cm^-1^.

## References

1. Brundage, L., J.P. Hendrick, E. Schiebel, a.J.M. Driessen, and W. Wickner. 1990. The purified E. coli integral membrane protein SecY/E is sufficient for reconstitution of SecA-dependent precursor protein translocation. Cell. 62 649–657.

2. Görlich, D., and T.A. Rapoport. 1993. Protein translocation into proteoliposomes reconstituted from purified components of the endoplasmic reticulum membrane. Cell. 75: 615–630.

3. Rapoport, T.A., L. Li, and E. Park. 2017. Structural and Mechanistic Insights into Protein Translocation. Annu. Rev. Cell Dev. Biol. 33: 369–390.

4. Cranford-Smith, T., and D. Huber. 2018. The way is the goal: how SecA transports proteins across the cytoplasmic membrane in bacteria. FEMS Microbiol. Lett. 365.

5. Pfeffer, S., L. Burbaum, P. Unverdorben, M. Pech, Y. Chen, R. Zimmermann, R. Beckmann, and F. Förster. 2015. Structure of the native Sec61 protein-conducting channel. Nat. Commun. 6: 8403.

6. Duong, F., and W. Wickner. 1997. The SecDFyajC domain of preprotein translocase controls preprotein movement by regulating SecA membrane cycling. EMBO J. 16: 4871–4879.

7. Scotti, P.A., M.L. Urbanus, J. Brunner, J.W. de Gier, G. von Heijne, C. van der Does, A.J. Driessen, B. Oudega, and J. Luirink. 2000. YidC, the Escherichia coli homologue of mitochondrial Oxa1p, is a component of the Sec translocase. EMBO J. 19: 542–9.

8. Schulze, R.J., J. Komar, M. Botte, W.J. Allen, S. Whitehouse, V.A.M. Gold, J.A. Lycklama, A. Nijeholt, K. Huard, I. Berger, C. Schaffitzel, and I. Collinson. 2014.Membrane protein insertion and proton-motive-force-dependent secretion through the bacterial holo-translocon SecYEG-SecDF-YajC-YidC. Proc. Natl. Acad. Sci. U. S. A. 111: 4844–9.

9. Müller, M., H.G. Koch, K. Beck, and U. Schäfer. 2001. Protein traffic in bacteria: multiple routes from the ribosome to and across the membrane. Prog. Nucleic Acid Res. Mol. Biol. 66: 107–57.

10. Bieniossek, C., Y. Nie, D. Frey, N. Olieric, C. Schaffitzel, I. Collinson, C. Romier, P. Berger, T.J. Richmond, M.O. Steinmetz, and I. Berger. 2009. Automated unrestricted multigene recombineering for multiprotein complex production. Nat. Methods. 6: 447–450.

11. Komar, J., S. Alvira, R. Schulze, R. Martin, J. Lycklama a Nijeholt, S. Lee, T. Dafforn, G. Deckers-Hebestreit, I. Berger, C. Schaffitzel, and I. Collinson. 2016. Membrane protein insertion and assembly by the bacterial holo-translocon SecYEG-SecDF-YajC-YidC. Biochem. J. 0: 1–35.

12. Botte, M., N.R. Zaccai, J.L.à. Nijeholt, R. Martin, K. Knoops, G. Papai, J. Zou, A. Deniaud, M. Karuppasamy, Q. Jiang, A.S. Roy, K. Schulten, P. Schultz, J. Rappsilber, G. Zaccai, I. Berger, I. Collinson, and C. Schaffitzel. 2016. A central cavity within the holo-translocon suggests a mechanism for membrane protein insertion. Sci. Rep. 6: 38399.

13. Xu, Z., a L. Horwich, and P.B. Sigler. 1997. The crystal structure of the asymmetric GroEL-GroES-(ADP)7 chaperonin complex. Nature. 388: 741–750

14. Van Den Berg, B., W.M.C. Jr, I. Collinson, Y. Modis, E. Hartmann, S.C. Harrison, T.A. Rapoport, B. Van den Berg, and W.M. Clemons. 2004. X-ray structure of a protein-conducting channel. Nature. 427: 36–44.

15. Kumazaki, K., S. Chiba, M. Takemoto, A. Furukawa, K. Nishiyama, Y. Sugano, T. Mori, N. Dohmae, K. Hirata, Y. Nakada-Nakura, A.D. Maturana, Y. Tanaka, H. Mori, Y. Sugita, F. Arisaka, K. Ito, R. Ishitani, T. Tsukazaki, and O. Nureki. 2014. Structural basis of Sec-independent membrane protein insertion by YidC. Nature. 509: 516–20.

16. Tsukazaki, T., H. Mori, Y. Echizen, R. Ishitani, S. Fukai, T. Tanaka, A. Perederina, D.G. Vassylyev, T. Kohno, A.D. Maturana, K. Ito, and O. Nureki. 2011. Structure and function of a membrane component SecDF that enhances protein export. Nature. 474: 235–8.

17. Midtgaard, S.R., T.A. Darwish, M.C. Pedersen, P. Huda, A.H. Larsen, G.V. Jensen, S.A.R. Kynde, N. Skar-Gislinge, A.J.Z. Nielsen, C. Olesen, M. Blaise, J.J. Dorosz, T.S. Thorsen, R. Venskutonytė, C. Krintel, J. V. Møller, H. Frielinghaus, E.P. Gilbert, A. Martel, J.S. Kastrup, P.E. Jensen, P. Nissen, and L. Arleth. 2018. Invisible detergents for structure determination of membrane proteins by small-angle neutron scattering. FEBS J. 285: 357–371.

18. Pedersen, M.C., L. Arleth, and K. Mortensen. 2013. WillItFit: A framework for fitting of constrained models to small-angle scattering data. J. Appl.Crystallogr.

19. Persson, F., P. Söderhjelm, and B. Halle. 2018. The geometry of proteinhydration. J. Chem. Phys. 148.

20. Lomize, M.A., I.D. Pogozheva, H. Joo, H.I. Mosberg, and A.L. Lomize. 2012. OPM database and PPM web server: Resources for positioning of proteins in membranes. Nucleic Acids Res. 40: 370–376.

21. Hansen, S. 2012. BayesApp: A web site for indirect transformation of small-angle scattering data. J. Appl. Crystallogr. 45: 566–567.

22. Armen, R.S., O.D. Uitto, and S.E. Feller. 1998. Phospholipid component volumes: Determination and application to bilayer structure calculations. Biophys. J. 75: 734–744.

23. Larsen, A.H., J. Dorosz, T.S. Thorsen, N.T. Johansen, T. Darwish, S.R. Midtgaard, L. Arleth, and J.S. Kastrup. 2018. Small-angle neutron scattering studies on the AMPA receptor GluA2 in the resting, AMPA-bound and GYKI-53655-bound states. IUCrJ. 5: 1–14.

24. Teixeira, J. 1988. Small-angle scattering by fractal systems. J. Appl. Crystallogr. 21: 781–785.

25. Kotlarchyk, M., and S. Chen. 1983. Analysis of small angle neutron scattering spectra from polydisperse interacting colloids. J. Chem. Phys. 79: 2461.

26. Svergun, D., C. Barberato, and M.H. Koch. 1995. CRYSOL - A program to evaluate X-ray solution scattering of biological macromolecules from atomic coordinates. J. Appl. Crystallogr. 28: 768–773.

27. Stansfeld, P.J., J.E. Goose, M. Caffrey, E.P. Carpenter, J.L. Parker, S. Newstead, and M.S.P. Sansom. 2015. MemProtMD: Automated Insertion of Membrane Protein Structures into Explicit Lipid Membranes. Structure. 23: 1350–1361.

28. Stansfeld, P.J., and M.S.P. Sansom. 2011. From coarse grained to atomistic: A serial multiscale approach to membrane protein simulations. J. Chem. Theory Comput. 7: 1157–1166.

29. Guinier, A., and G. Fournet. 1955. Small-angle scattering of X-rays. Wiley.

30. Furukawa, A., K. Yoshikaie, T. Mori, H. Mori, Y.V. Morimoto, Y. Sugano, S. Iwaki, T. Minamino, Y. Sugita, Y. Tanaka, and T. Tsukazaki. 2017. Tunnel Formation Inferred from the I-Form Structures of the Proton-Driven Protein Secretion Motor SecDF. Cell Rep. 19: 895–901.

31. Larsen, A.H., L. Arleth, S. Hansen, and IUCr. 2018. Analysis of small-angle scattering data using model fitting and Bayesian regularization. J. Appl. Crystallogr. 51: 1151–1161.

32. Kumazaki, K., T. Kishimoto, A. Furukawa, H. Mori, Y. Tanaka, N. Dohmae, R. Ishitani, T. Tsukazaki, and O. Nureki. 2015. Crystal structure of Escherichia coli YidC, a membrane protein chaperone and insertase. Sci. Rep. 4: 7299.

33. Tanaka, Y., Y. Sugano, M. Takemoto, T. Mori, A. Furukawa, T. Kusakizako, K. Kumazaki, A. Kashima, R. Ishitani, Y. Sugita, O. Nureki, and T. Tsukazaki.2015. Crystal Structures of SecYEG in Lipidic Cubic Phase Elucidate a Precise Resting and a Peptide-Bound State. Cell Rep. 13: 1561–1568.

34. Eswar, N., B. Webb, M.A. Marti-Renom, M.S. Madhusudhan, D. Eramian, M. Shen, U. Pieper, and A. Sali. 2007. Comparative Protein Structure Modeling Using MODELLER. In: Current Protocols in Protein Science. Hoboken, NJ, USA: John Wiley & Sons, Inc. p. 2.9.1–2.9.31.

35. Marrink, S.J., H.J. Risselada, S. Yefimov, D.P. Tieleman, and A.H. De Vries. 2007. The MARTINI force field: Coarse grained model for biomolecular simulations. J. Phys. Chem. B. 111: 7812–7824.

36. Monticelli, L., S.K. Kandasamy, X. Periole, R.G. Larson, D.P. Tieleman, and S.J. Marrink. 2008. The MARTINI coarse-grained force field: Extension to proteins. J. Chem. Theory Comput. 4: 819–834.

37. Bechara, C., A. Nöll, N. Morgner, M.T. Degiacomi, R. Tampé, and C. V. Robinson. 2015. A subset of annular lipids is linked to the flippase activity of an ABC transporter. Nat. Chem. 7: 255–262.

38. Laganowsky, A., E. Reading, T.M. Allison, M.B. Ulmschneider, M.T. Degiacomi, A.J. Baldwin, and C. V. Robinson. 2014. Membrane proteins bind lipids selectively to modulate their structure and function. Nature. 510: 172–175.

39. Corey, R.A., E. Pyle, W.J. Allen, D.W. Watkins, M. Casiraghi, B. Miroux, I. Arechaga, A. Politis, and I. Collinson. 2018. Specific cardiolipin-SecY interactions are required for proton-motive force stimulation of protein secretion. Proc. Natl. Acad. Sci. 115: 7967–7972.

40. Nagamori, S., I.N. Smirnova, and H.R. Kaback. 2004. Role of YidC in folding of polytopic membrane proteins. J. Cell Biol. 165: 53–62.

41. Zhu, L., H.R. Kaback, and R.E. Dalbey. 2013. YidC protein, a molecular chaperone for LacY protein folding via the SecYEG protein machinery. J. Biol. Chem. 288: 28180–28194.

